# A thalamo-centric neural signature for restructuring negative self-beliefs

**DOI:** 10.1101/2021.08.26.457858

**Authors:** Trevor Steward, Po-Han Kung, Christopher G. Davey, Bradford A. Moffat, Rebecca K. Glarin, Alec J. Jamieson, Kim L. Felmingham, Ben J. Harrison

**Affiliations:** Melbourne Neuropsychiatry Centre, Department of Psychiatry, The University of Melbourne, Victoria, Australia; Melbourne School of Psychological Sciences, Faculty of Medicine, Dentistry and Health Sciences, University of Melbourne, Parkville, Victoria, Australia; The Melbourne Brain Centre Imaging Unit, Department of Medicine and Radiology, The University of Melbourne

## Abstract

Negative self-beliefs are a core feature of psychopathology. Despite this, we have a limited understanding of the brain mechanisms by which negative self-beliefs are cognitively restructured. Using a novel paradigm, we had participants use Socratic questioning techniques to restructure self-beliefs during ultra-high resolution 7-Tesla functional magnetic resonance imaging (UHF fMRI) scanning. Cognitive restructuring elicited prominent activation in a fronto-striato-thalamic circuit, including the mediodorsal thalamus (MD), a group of deep subcortical nuclei believed to synchronize and integrate prefrontal cortex activity, but which has seldom been directly examined with fMRI due to its small size. Increased activity was also identified in the medial prefrontal cortex (MPFC), a region consistently activated by internally focused mental processing, as well as in lateral prefrontal regions associated with regulating emotional reactivity. Using Dynamic Causal Modelling (DCM), evidence was found to support the MD as having a strong excitatory effect on the activity of regions within the broader network mediating cognitive restructuring. Moreover, the degree to which participants modulated MPFC-to-MD effective connectivity during cognitive restructuring predicted their individual tendency to engage in repetitive negative thinking. Our findings represent a major shift from a cortico-centric framework of cognition and provide important mechanistic insights into how the MD facilitates key processes in cognitive interventions for common psychiatric disorders. In addition to relaying integrative information across basal ganglia and the cortex, we propose a multifaceted role for the MD whose broad excitatory pathways act to increase synchrony between cortical regions to sustain complex mental representations, including the self.

## Introduction

Beliefs that are negatively biased, inaccurate, and rigid play a key role in etiology and maintenance of psychopathology [1]. Cognitive models of mood disorders posit that maladaptive self-beliefs – for example, believing that one is inherently flawed or unlovable – are central to triggering the emotional disturbances characteristic of these disorders [2]. Cognitive-behavioral therapy (CBT) and other evidence-based psychotherapeutic treatments are centered on identifying and restructuring maladaptive cognitions, often through Socratic questioning techniques [3, 4]. The extent to which an individual’s self-beliefs are malleable has been found to uniquely predict CBT outcomes, as well as long-term reductions in disorder severity [5]. Despite the importance of cognitive restructuring in psychotherapeutic interventions, the neurobiological mechanisms underpinning these processes remain largely enigmatic.

Neuroimaging studies have identified a consistent set of brain regions that support the cognitive reappraisal of negative emotions elicited by provocative stimuli, typically visual images. This network comprises the dorsolateral prefrontal cortex (dlPFC), pre-supplementary motor area (preSMA) and the dorsal anterior cingulate cortex (dACC) [6– 8], whose increased activity consistently accompanies the successful down-regulation of emotion, together with the modulation of activity in other brain regions, including the amygdala [9, 10]. Whether this network also supports the cognitive restructuring of negative self-beliefs remains unclear. In a recent study of patients with social anxiety disorder, a more distinct involvement of the ‘default mode network (DMN)’ was reported when patients reacted to versus accepted negative self-beliefs [11]. This finding is broadly consistent with other studies linking DMN activity, particularly the medial prefrontal cortex (MPFC), to negative self-appraisal processes including depressive rumination [12, 13]. Based on such findings, it has been hypothesized that this core MPFC-based ‘self-network’ dynamically interacts with lateral PFC ‘control’ regions and affective value-signaling regions, including the striatum [14, 15]. Within this framework, subcortical network hubs are likely to be pivotal for integrating self-representations across distributed cortical regions in support of higher-order cognitive processing [16, 17].

Of these subcortical regions, the mediodorsal thalamus (MD) may be especially important to higher-order cognition due to its distinguishing arrangement of dense direct and converging innervations from multiple PFC regions [18]. These features suggest that the MD serves as a relay that rapidly enables crosstalk between widely distributed PFC regions – a major shift from the ‘cortico-centric’ view of cognition [19]. Lesions to the MD induce cognitive abnormalities similar to those observed with focal PFC damage [20], indicating that higher-order cognition cannot be fully understood without examining the reciprocal thalamocortical circuitry that underpins it. Furthermore, complementing its close partnership with the PFC, the MD receives dense afferent projections from basal ganglia structures, including the striatum [21]. In a noteworthy recent study, multiple large-scale cortical networks, spanning the DMN and lateral PFC, were shown to converge on primary subcortical connectivity zones in the MD and caudate nucleus (CN) – suggesting a putative mechanism for the dynamic integration of cognitive networks supporting aforementioned self and control processes [22]. Taken together, the divergent and convergent nature of MD projections offers a plausible architecture for integrating multiple information streams, thereby enabling the elaboration of complex mental representations required for cognitive restructuring [23].

Our aim in the current study was to investigate the neural systems basis of restructuring negative self-beliefs with an emphasis on mapping integrative fronto-striato-thalamic circuit activity. To do so, we developed a novel regulation paradigm in which participants were trained to cognitively restructure negative self-beliefs using established Socratic questioning techniques (Fig. 1). To support the precise functional anatomical mapping of fronto-striato-thalamic regions, we utilized ultra-high field functional magnetic resonance imaging (UHF fMRI), which we combined with dynamic causal modelling (DCM) to examine directed interactions (‘effective connectivity’) between identified task-responsive regions. Our principal hypothesis was that the cognitive restructuring of negative beliefs would elicit robust activation of a fronto-striato-thalamic network, encompassing the MD and CN as major subcortical hubs, together with the MPFC, dACC/preSMA and dlPFC. Through DCM, we tested multiple architectures of fronto-striato-thalamic function with the specific goal of mapping the causal influence of the MD on broader network activity. Lastly, we examined whether participants’ tendency to engage in repetitive negative thinking – a trait construct broadly linked to affective disorders [1]– could be predicted by fronto-striato-thalamic interactions during cognitive restructuring.

**Figure 1.**
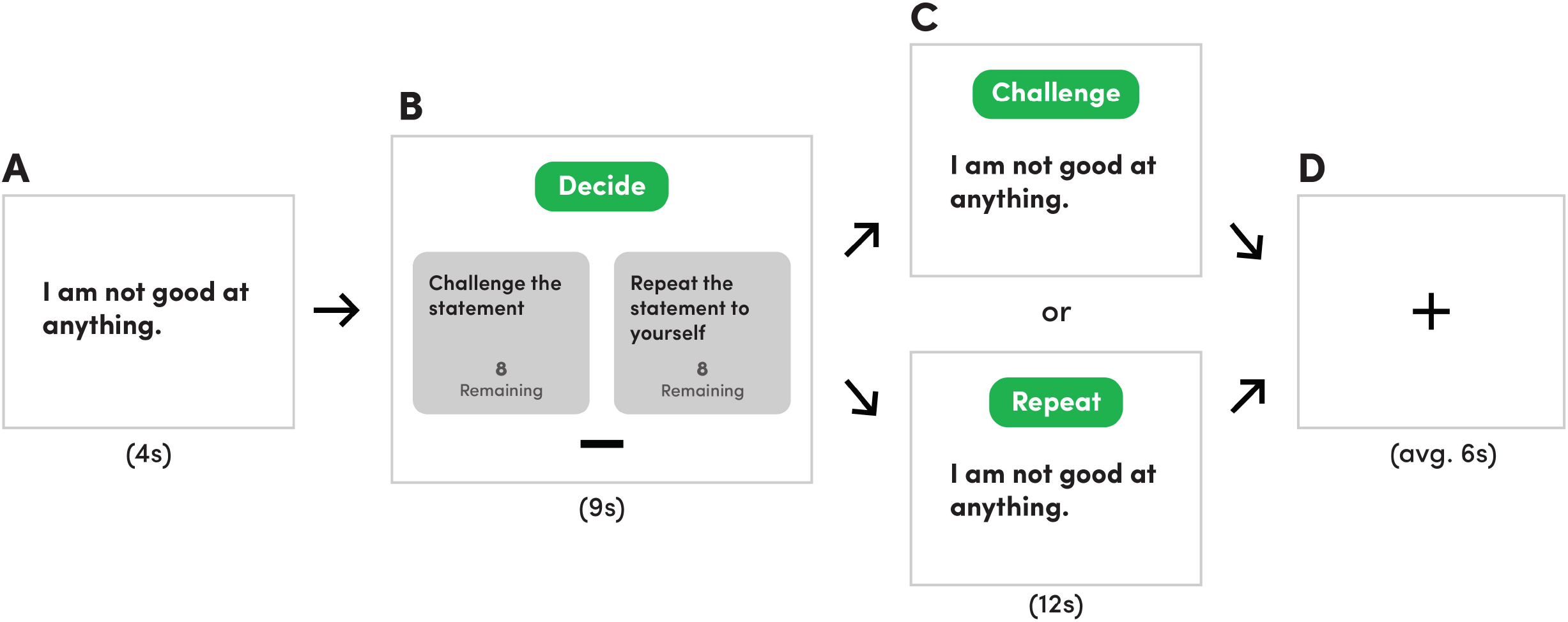
Cognitive restructuring paradigm. The task consisted of one run of 16 blocks. In each block, (A) a negative belief statement was presented for 4 seconds; (B) next, participants had 9 seconds to decide whether to challenge (CHAL) the statement using previously trained Socratic questioning techniques or to repeat (REP) the statement to themselves. During each block, participants were shown how many remaining choices they had for each option (CHAL or REP); (C) after their choice, participants either cognitively reframed or repeated the statement for 12 seconds; (D) a fixation cross was displayed for an average of 6 seconds before the next block began.

## Materials and Methods

### Participants

49 healthy adults were recruited for this study. Exclusion criteria included: 1) the presence of a serious mental illness (e.g., psychosis, bipolar disorder, obsessive-compulsive disorder) screened using the Mini-International Neuropsychiatric Interview (MINI; [24]; or 2) MRI contraindications (e.g., pregnancy, metallic implants or claustrophobia). Participants attended a single testing session at the Melbourne Brain Centre Imaging Unit (The University of Melbourne, Parkville). This study was approved by The University of Melbourne Human Research Ethics Committee. Seven participants were excluded from this initial sample due to poor physiological recording quality (n=3), not properly completing the task (n=2), scanning being terminated early (n=1), and excessive movement (n=1). As a result, 42 healthy participants (age 24.7 ± 4.65 years) were included in the final DCM analysis. Table S1 provides a summary of the sociodemographic characteristics of the final sample.

### Behavioral Measures

Repetitive negative thinking was measured using the Perseverative Thinking Questionnaire (PTQ). The validity is of this questionnaire is supported by associations with existing repetitive negative thinking measures and by correlations with depression and anxiety symptom levels [25]. Internal consistency for the Perseverative Thinking Questionnaire (PTQ) was excellent (Cronbach’s α = 0.908). All behavioral and connectivity data were checked for normality of distribution by Shapiro–Wilk tests before performing parametric statistical analyses.

### MRI Paradigm

Further information on the MRI paradigm training procedures is available in the Supplementary Information.

The cognitive-restructuring task consisted of a single run containing sixteen blocks (see Fig. 1). In each block, participants were first presented with a negative self-belief statement for four seconds. Next, participants were given nine seconds to decide whether to challenge (i.e., cognitively reframe) or repeat the statement. Participants indicated their choices using an MRI-compatible control pad. To ensure sufficient an equal number of blocks for each condition, participants were instructed to only cognitively restructure half the statements and to repeat the remaining half of the statements. The remaining number of times that participants could challenge or repeat a statement was displayed alongside these two choices. After their choice, the same statement was displayed for twelve seconds, during which time participants engaged in their previously selected strategy. If the participant chose to restructure the negative belief, a prompt reading ‘Challenge’ was displayed alongside the statement, and participants were instructed to mentally refute or reinterpret the negative statement throughout the entire twelve seconds (herein referred to as ‘CHAL’). If the choice were to repeat, a prompt reading ‘Repeat’ was shown with the negative statement, and participants were instructed to mentally recite the negative belief until the twelve seconds had expired (herein referred to as ‘REP’). Between each statement block, a fixation cross was presented for an average of six seconds to reduce carry-over effects.

### Image Acquisition

Information on 7-Tesla (7T) image acquisition parameters, preprocessing and physiological noise correction is available in the Supplementary Information.

### First- and second-level General Linear Model (GLM) Analyses

First-level (single-subject) contrast images were estimated for CHAL>REP to characterize changes in brain activation associated with cognitive restructuring. Participant’s pre-processed timeseries and nuisance regressors (i.e., physiological noise and motion fingerprint regressors) were included in the GLM analysis, with the onset times for each condition event specified and convolved with the SPM canonical hemodynamic response function (HRF). A 128-Hz high-pass filter was applied to account for low-frequency noise. Temporal autocorrelation was estimated using SPM’s FAST method, which has been shown to outperform AR(1) at short TRs and yield superior reliability [26]. Contrast images for each participant were entered in a second-level random-effects GLM using a one-sample t-test design. For all GLM analyses, whole-brain, false discovery rate (FDR) corrected statistical thresholds were applied (P_FDR_ <0.05), in addition to a 10-voxel cluster-extent threshold (K_E_≥10 voxels).

### Dynamic Causal Modelling (DCM)

DCM estimates the directional interactions between brain regions of interest through the process of generating timeseries from underlying neurobiological causes. These timeseries are dependent upon the connectivity architecture of the network and the strength of the connectivity parameters. Parameter strengths are estimated through model inversion, a process of finding the parameters that offer the best trade-off between model fit and model complexity. The relative evidence of these estimates can then be compared through model comparison, which tests hypothetical functional architectures to identify a model which optimally explains the data. In contrast to temporal correlation measures of functional connectivity, DCM provides a more detailed and physiologically valid mapping of effective connectivity – the directed causal influences of brain regions on one another [27]. In DCM, modulation is measured in hertz (Hz), which denotes the rates of change in activity caused by the dynamic influence of one region on another. Positive effective connectivity indicates a putative excitatory upregulation of activity, whereas negative connectivity represents an inhibitory downregulation by a task effect (i.e., cognitive restructuring). It has been shown that 7T fMRI furnishes more efficient estimates of effective connectivity than those provided by lower field strengths [28].

In addition to the left MD, peaks from four regions displaying significant changes in activation during the CHAL>REP contrast were included in the model space: the left CN, ventral MPFC, dlPFC and preSMA. The regional time-series (VOIs) for each of these areas were extracted at an individual subject level following recently published guidelines [29]. Further information on our VOI extraction procedure in the Supplementary Information.

Our full model space was specified using DCM 12.5. Models varied by two components: the endogenous connections between nodes (intrinsic parameters) and the modulation of the strengths of functional coupling between the MD and other regions induced by cognitive restructuring. The full model assumed bidirectional endogenous connections between the MD with the CN, ventral MPFC, dlPFC and the preSMA (Fig. 2B) and set ‘Task’, the onset of all blocks comprising both CHAL and REP blocks, as the driving input to all regions. In order to test the modulatory effect of cognitive restructuring on MD connectivity, CHAL was set as the modulatory input on each connection to and from the MD. BMR tested whether modulation by CHAL occurred on the connections from the CN, ventral MPFC, dlPFC, and preSMA to the MD, or from the MD to the CN, ventral MPFC, dlPFC, and preSMA.

**Figure 2.**
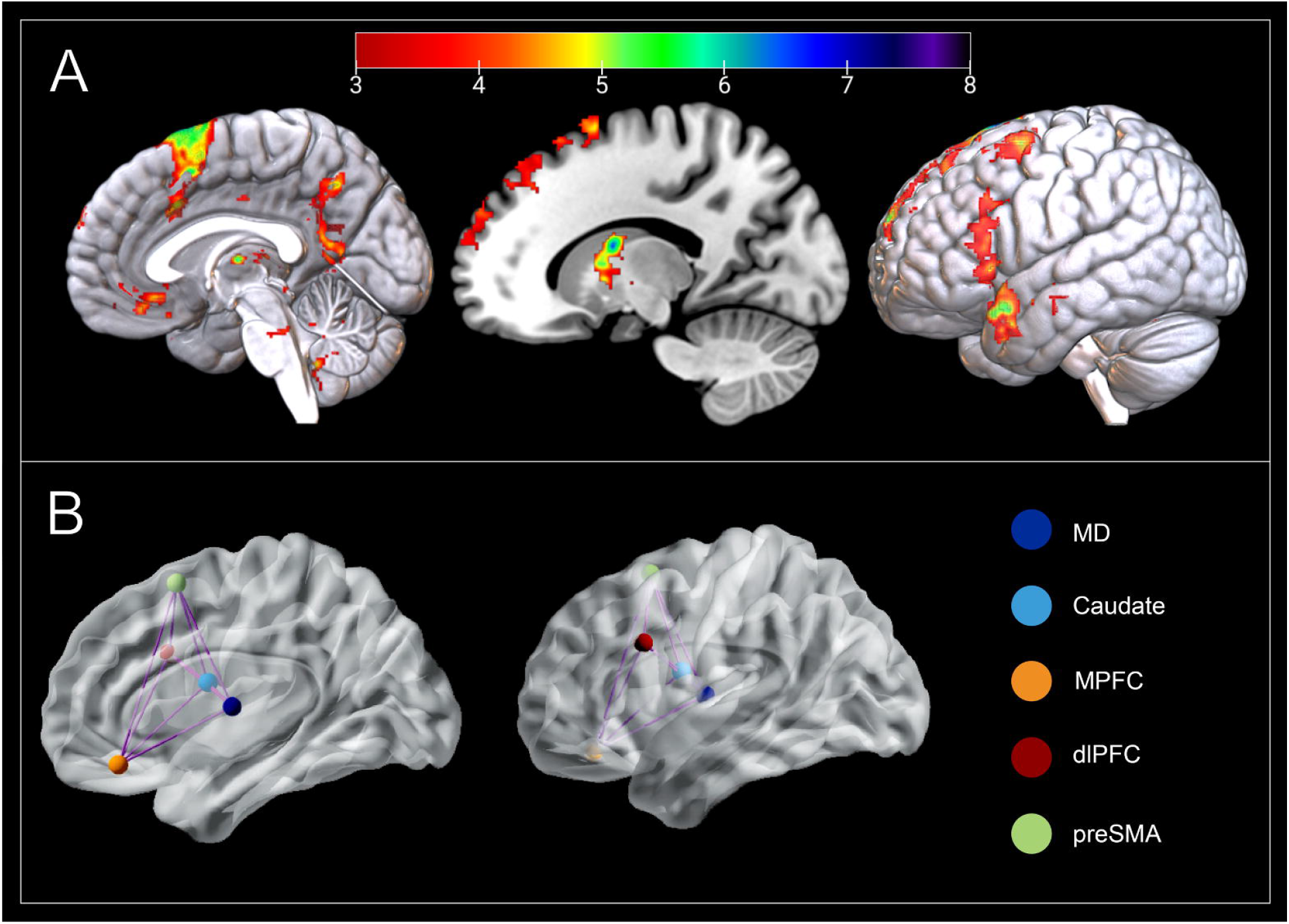
Neural response elicited by cognitive restructuring. (A) The mediodorsal thalamus (MD; x = -2; y = -10; z = 8), caudate (x = -16; y = 0; z = 18), medial prefrontal cortex (MPFC; x = -2; y = 37; z = -16), dorsolateral prefrontal cortex (dlPFC; x= -43; y = 16; z = 30), and pre-supplementary motor area (pre-SMA; x = -8; y = 11; z = 67) are responsive to cognitive restructuring (CHAL>REP). Group activation is overlaid on a MNI T1 template. Color bar represents t-values. (B) Our full model assumed bidirectional intrinsic connectivity between all five DCM nodes.

Full models of effective connectivity were fitted to each participant’s timeseries data, yielding posterior connectivity parameters and their probabilities. At the group level, the posterior connectivity parameter estimates from all participants’ DCMs were assessed using Parametric Empirical Bayes (PEB) and Bayesian Model Reduction (BMR). The PEB framework affords robust group-level analyses of effective connectivity by means of a hierarchical model, comprising DCMs at the single-subject level and a GLM of connectivity parameters between subjects [30]. After estimating the PEB model, parameters that did not contribute to the model evidence were pruned using BMR. Posterior parameter estimates following BMR were averaged using Bayesian model averaging (BMA), and the ensuing BMA parameters (with a posterior probability > 95%) are reported in the Supplementary information, Figure S2. The resulting pattern of effective connectivity is illustrated in Fig. 3.

**Figure 3.**
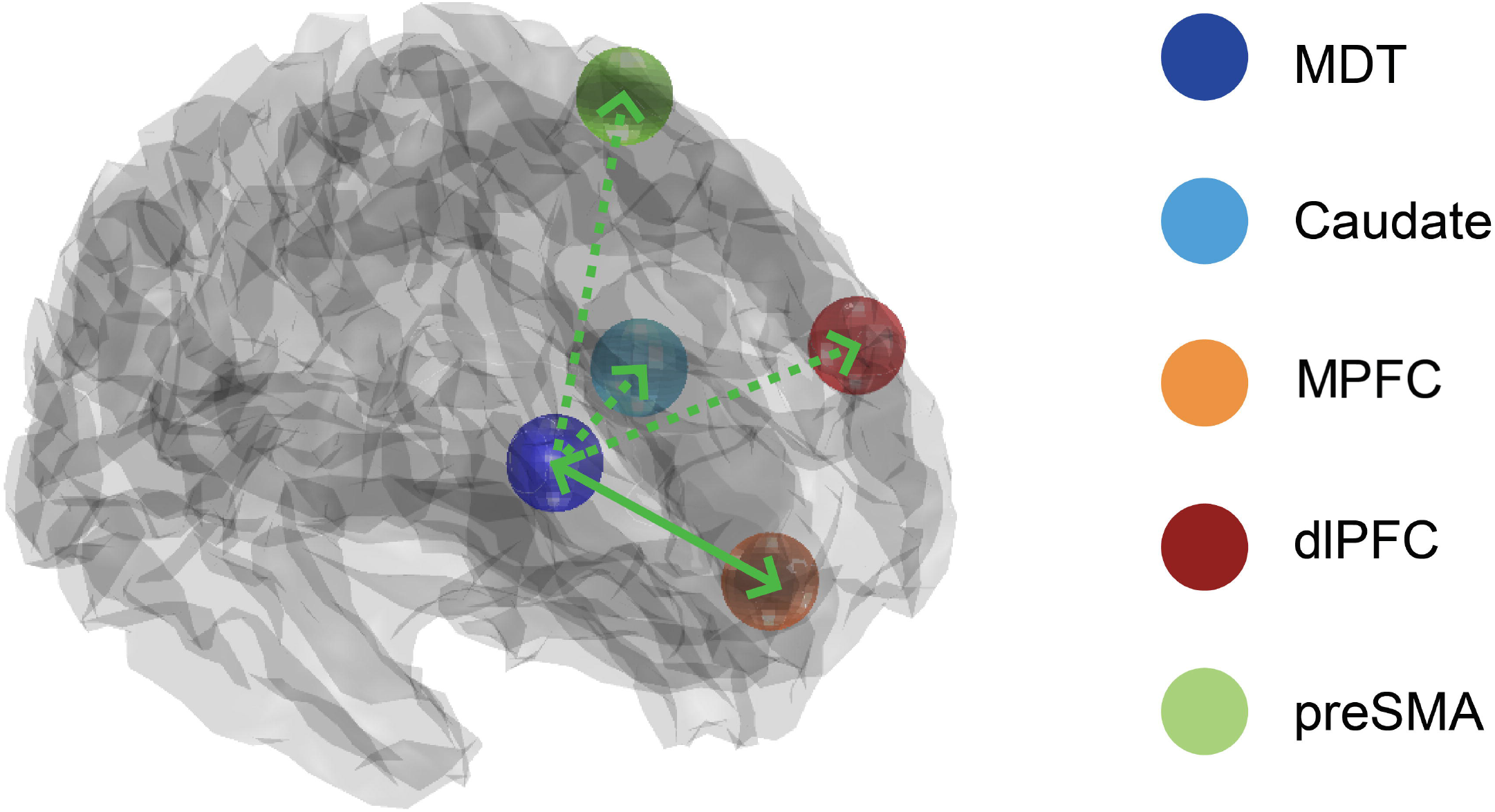
Mediodorsal thalamus effective connectivity and its relationship to cognitive restructuring. Carrying out cognitive restructuring led to excitatory effects on effective connectivity from the mediodorsal thalamus (MD) to the caudate, medial prefrontal cortex (MPFC), dorsolateral prefrontal cortex (dlPFC), and pre-supplementary motor area (pre-SMA; posterior probability > .95). In contrast, evidence was only found for excitatory effects from the ventral MPFC to the MD during cognitive restructuring (posterior probability > .95). Unilateral modulation of pathways from the MD is depicted with a broken green line. Bilateral modulations of MD pathways are depicted with a solid green.

Last, we sought to test whether the effect of modulation of MD pathways by cognitive restructuring depended on individual participants’ tendency to engage in repetitive negative thinking. A PEB model containing a regressor quantifying the effects of PTQ scores on each of DCM modulatory parameter was examined, with parameters which did not contribute to model evidence being removed through pruning. Parameters featuring strong evidence (posterior probability > 95%) were then tested using leave one-out cross validation to determine whether the size of these parameters were sufficiently large to predict a participant’s PTQ score. This procedure iteratively creates a PEB model on all expect for one subject, then predicts the PTQ score of the left-out subject. These complete results are available at https://github.com/trevorcsteward/MD_Effective_Connectivity.

## Results

### Behavioral Results

After extensive training on how to use Socratic questioning to challenge negative self-beliefs, participants completed our cognitive restructuring task during which they were presented with common negative self-belief statements during scanning (see Fig. 1 and Materials and Methods). We observed a significant decrease from pre- to post-scanning in participants’ endorsement of the negative self-beliefs statements that were restructured (50% of statements; t(48) = 7.85, p < .001), indicating that they successfully applied Socratic questioning to restructure negative self-beliefs.

### GLM Results

As hypothesized, whole-brain fMRI analysis (p<0.05, voxelwise false discovery rate [FDR] corrected) confirmed that cognitive restructuring, compared to the task repeat condition, elicited significant activation of distributed frontal cortical and striatal-thalamic regions (Fig. 2A). Cortically, these regions included the MPFC, specifically its ventral-subgenual aspect; the preSMA extending to dACC, as well as the left dlPFC and frontal operculum. Subcortically, we observed prominent activation of the MD and head of the CN, together with other basal ganglia regions including the globus pallidus and ventral putamen, as well as the midbrain periaqueductal gray. A complete anatomical description of these results is provided in Supplementary Table S2.

### DCM Results

DCM utilizes a Bayesian framework to infer the causal architecture of a network of regions (i.e., nodes), defined in terms their ‘effective connectivity’ – the extent to which a region’s activity directly influences another. Here, we used DCM used to assess the modulatory impact of cognitive restructuring on MD effective connectivity with other key regions of interest, including the ventral MPFC, dlPFC, preSMA and CN (Fig. 2B). The full model of our network was designed to examine the modulatory effects of cognitive restructuring on pathways to- and from the MD. Bayesian model reduction (BMR) was used to iteratively test configurations of this neural architecture and to prune any redundant parameters which did not contribute to the evidence. This method allows for the identification regional connectivity parameters best explained by the data at a group level and for the verification of causal excitatory or inhibitory effects of MD circuits of brain activity during cognitive restructuring [31].

Consistent with previous work postulating an influence of the MD on cortical systems [18], BMR identified a strong excitatory effect of MD activity on the ventral MPFC, dlPFC and preSMA during cognitive restructuring (posterior probability > 0.95; Fig. 3). MD activity also had an excitatory influence on the CN (head) during cognitive restructuring – a result that is consistent with evidence of direct anatomical connections between these regions [32]. Relevantly, the ventral MPFC was the only region identified to exert a reciprocal modulatory (excitatory) influence on MD activity, suggesting that this pathway may serve as the primary relay for PFC modulation of the MD across this network (Fig. 3 and Fig. S2).

### Mediodorsal Thalamus Modulation and Repetitive Negative Thinking

Using estimates of connectivity strengths, we tested whether MD interactions during cognitive restructuring predicted participants’ tendency to engage in repetitive negative thinking. A Parametric Empirical Bayes (PEB) model was specified containing a parameter to quantify the effects of individual Perseverative Thinking Questionnaire (PTQ) scores on each MD pathway in our network. There was strong evidence to support an association between higher PTQ scores and greater excitatory influence of the ventral MPFC on MD activity during cognitive restructuring (posterior probability > 0.95). Next, we used leave-one-out cross validation [31] to establish the predictive validity our model and found that individual ventral MPFC-to-MD connectivity strength levels could reliably classify participants’ PTQ scores (p=0.029, r=0.29; Fig. 4).

**Figure 4.**
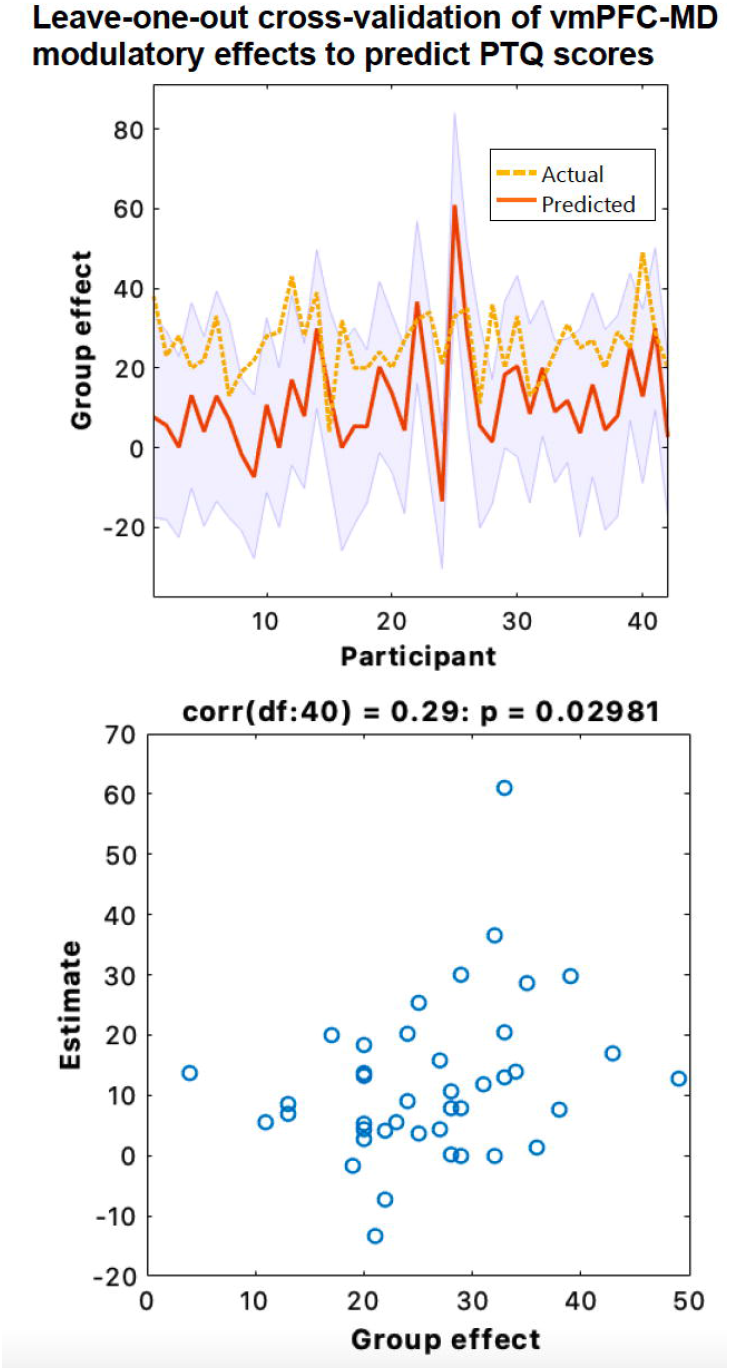
Leave-one-out cross validation of Parametric Empirical Bayes (PEB) effects on ventral MPFC-to-MD modulation during cognitive restructuring. Top panel: the out-of-samples estimate of Perseverative Thinking Questionnaire (PTQ) scores for each subject (red line) with 90% confidence interval (shaded area). The dashed orange line is the actual group effect. Bottom: The actual subject effect plotted against the expected value of the estimated subject effect. Participant PTQ scores can be reliably predicted based on their modulation of ventral MPFC-to-MD effective connectivity during cognitive restructuring (p=0.029, r=0.29)

## Discussion

Our aim was to assess how the cognitive restructuring of negative self-beliefs is mediated by fronto-striato-thalamic pathways, with particular emphasis on the functionally integrative role of the mediodorsal thalamus (MD). As hypothesized, cognitive restructuring elicited prominent activation of DMN regions associated with self-directed thought, lateral PFC cognitive control regions, and subcortical hubs including the MD and head of the CN. Using DCM, applied to UHF fMRI data, we found strong evidence for the MD having an excitatory effect on the PFC and CN during cognitive restructuring, as well as the MPFC demonstrating a reciprocal excitatory effect on the MD. Together, our findings endorse the MD as having a central role in mediating higher-order cognition by integrating and sustaining activity across widespread frontal regions. This study also provides important knowledge into the mechanistic changes that neural networks may undergo during psychotherapeutic interventions like cognitive therapy.

Neuroimaging research has provided key insights into the neural systems underpinning the regulation of affective and self-referential processing [15]. Based upon previous evidence demonstrating that the DMN is reliably activated when reflecting on one’s own traits and autobiographical knowledge [12, 13, 33, 34], we correctly hypothesized that the cognitive reframing of negative self-beliefs would elicit activity in anterior DMN regions, including the MPFC. It has been posited that the DMN contributes to the representation of the self as object, with MPFC subregions acting to select and gate these representations into conscious awareness [14]. By bringing conceptual and associative knowledge to bear on current thought and perception, this processing stream is ideally positioned to oversee introspective processes and to regulate self-referential thoughts [35]. Restructuring negative self-beliefs was also found to result in broad activation of lateral PFC control regions and anterior midline regions, namely the preSMA and dACC, which is broadly consistent with prior studies of cognitive emotion regulation [7, 9, 11]. Instead of observing suppression of DMN function, which is frequent in cognitively demanding tasks [36], restructuring negative self-beliefs evoked its joint engagement with valuation and regulatory control networks. As a whole, these effects support this novel paradigm recruiting a unique self-directed cognition whose synergistic interactions facilitate the regulation of self-representations in a constructive manner in order to enable the restructuring of negative self-beliefs.

We identified heightened activity during cognitive restructuring in subcortical structures, namely the MD, that have recently been found to form part of a more comprehensive neuroanatomical model of the DMN [37]. Extensive research from the past two decades has established a coordinating role for the MD in distinct cognitive domains that are mediated by PFC regions [18, 19, 23]. However, it has been unclear to what degree do parallel fronto-striato-thalamic circuits underlie higher-order cognitive functions, such as processing reflective cortical representations of the self. Our findings demonstrating strong excitatory effects from the MD on the MPFC, dlPFC and preSMA during cognitive restructuring directly supports recent animal work showing that the MD amplifies and sustains local PFC connectivity to enable neural sequences to emerge that maintain mental representations [38]. We posit that, rather than solely serving as a simple relay for PFC regions, excitatory MD pathways acts to increase PFC information convergence by recruiting previously untuned cortical neurons and increasing interregional synchrony, thus contributing to the generation of complex mental representations [19].

Although interactions between the thalamus and cortex are essential for cognition, there is increasing evidence that the MD may also have a role in sustaining and transmitting information on context-relevant representations to subcortical regions [23]. Our data was best explained by a model in which the MD had an excitatory effect on the CN during cognitive restructuring, indicating that the MD may transmit updated information to the basal ganglia in support of flexible goal-directed actions. Other animal studies have demonstrated that the inhibition of MD-striatal pathways prevents the incorporation information on internal states to guide decision-making [39]. Within the context of higher-order metacognition, the MD may therefore serve as a principal integrative nexus within fronto-striatal-thalamic loops, which acts to sustain and update the value assigned to mental representations. Our model supports such an update function, whereby the MD may receive *a priori* predictions as input from the cortex and subsequently projects *a posteriori* outcomes to other regions, including those in the basal ganglia [19, 40].

MD neurons are uniquely positioned in that they are capable of representing aggregations of cortical signals [41]. Varying levels of input convergence from the PFC, both in terms of magnitude and type, endow MD circuits with a computational role that is capable of simultaneously transforming multiple modalities of information [16, 42]. Findings from our DCM analysis endorse the ventral MPFC as the primary conduit for prefrontal input onto the MD during higher-order cognition. This result aligns with recent in-vitro electrophysiological research identifying distinct roles of prefrontal-MD pathways in shaping behavior, with inhibition of ventral MPFC-to-MD pathways impacting the encoding and maintenance of contingencies [43]. The fact that the modulation of ventral MPFC-to-MD effective connectivity during cognitive restructuring was able to reliably predict participants’ tendency to engage in repetitive negative thinking suggests that this pathway may contribute to sustaining mental self-representations and, when having exaggerated impact, to rumination. That is, individuals who excessively recruit this pathway may be less disposed to cognitively reframing maladaptive negative self-beliefs and more prone to engaging in repetitive negative thinking or rumination. Given the fundamental role of negative self-beliefs in multiple forms of psychopathology [4], future longitudinal studies using similar paradigms may have translational relevance in assessing the malleability of neural networks underlying self-beliefs.

## Conclusion

Our UHF fMRI study demonstrates that cognitively restructuring negative self-beliefs is supported by a distinct fronto-striato-thalamic circuitry, consistent with recent models of the self as a complex dynamic entity that emerges from the coordinated activity of multiple interacting brain systems [12–15]. Within this unique circuitry, we have confirmed for the first time a key excitatory role for the MD in humans [44], in which MD amplification acts to increase activity in multiple cortical regions during higher-order processes, as well as for the MPFC having a reciprocal excitatory effect on MD activity. Moreover, our observed relationship between MPFC-MD connectivity and individual differences in repetitive negative thinking suggests that MD pathways may represent a potential focal stimulation target for common mental health disorders [45]. Taken together, these findings advance a multifaceted framework for the MD in which it acts to increase synchrony between cortical regions to enable the generation of complex self-representations required for cognitive restructuring.

## Acknowledgments

This study was supported by a National Health and Medical Research Council of Australia (NHMRC) Project Grant (1161897) to BJH. Trevor Steward is supported by a NHMRC/MRFF Investigator Grant (MRF1193736), a BBRF Young Investigator Grant, and a University of Melbourne McKenzie Fellowship. CGD was supported by an NHMRC Career Development Fellowship (1141738). BJH was supported by a NHMRC Career Development Fellowship (1124472). The authors thank Cristian Stella and Lisa Incerti for their contributions to data collection, and the participants for their involvement in the study. We acknowledge the facilities, and the scientific and technical assistance of the Australian National Imaging Facility, a National Collaborative Research Infrastructure Strategy (NCRIS) capability, at the Melbourne Brain Centre Imaging Unit (MBCIU), The University of Melbourne. The multiband fMRI sequence was generously supported by a research collaboration agreement with CMRR, University of Minnesota and the MP2RAGE works in progress sequence was provided by Siemens Healthineers (Germany) as advanced works in progress package.

## Data Availability

The code for the effective connectivity analyses is available within the SPM12 software package (https://www.fil.ion.ucl.ac.uk/spm). Effective connectivity data and the code used to generate the results presented here are available at https://github.com/trevorcsteward/MD_Effective_Connectivity.

## References

1. Ehring T, Watkins ER. Repetitive negative thinking as a transdiagnostic process. International Journal of Cognitive Therapy. 2008;1.

2. Beck AT, Dozois DJA. Cognitive therapy: Current status and future directions. Annual Review of Medicine. 2011;62.

3. Braun JD, Strunk DR, Sasso KE, Cooper AA. Therapist use of Socratic questioning predicts session-to-session symptom change in cognitive therapy for depression. Behaviour Research and Therapy. 2015;70.

4. Hofmann SG, Asmundson GJG, Beck AT. The Science of Cognitive Therapy. Behavior Therapy. 2013;44.

5. de Castella K, Goldin P, Jazaieri H, Heimberg RG, Dweck CS, Gross JJ. Emotion Beliefs and Cognitive Behavioural Therapy for Social Anxiety Disorder. Cognitive Behaviour Therapy. 2015;44.

6. Kohn N, Eickhoff SB, Scheller M, Laird AR, Fox PT, Habel U. Neural network of cognitive emotion regulation - An ALE meta-analysis and MACM analysis. NeuroImage. 2014;87.

7. Buhle JT, Silvers JA, Wage TD, Lopez R, Onyemekwu C, Kober H, et al. Cognitive reappraisal of emotion: A meta-analysis of human neuroimaging studies. Cerebral Cortex. 2014;24.

8. Morawetz C, Riedel MC, Salo T, Berboth S, Eickhoff SB, Laird AR, et al. Multiple large-scale neural networks underlying emotion regulation. Neuroscience and Biobehavioral Reviews. 2020;116.

9. Steward T, Davey CG, Jamieson AJ, Stephanou K, Soriano-Mas C, Felmingham KL, et al. Dynamic Neural Interactions Supporting the Cognitive Reappraisal of Emotion. Cerebral Cortex. 2021;31.

10. Berboth S, Morawetz C. Amygdala-prefrontal connectivity during emotion regulation: A meta-analysis of psychophysiological interactions. Neuropsychologia. 2021;153.

11. Dixon ML, Moodie CA, Goldin PR, Farb N, Heimberg RG, Gross JJ. Emotion Regulation in Social Anxiety Disorder: Reappraisal and Acceptance of Negative Self-beliefs. Biological Psychiatry: Cognitive Neuroscience and Neuroimaging. 2020;5.

12. Davey CG, Harrison BJ. The brain’s center of gravity: how the default mode network helps us to understand the self. World Psychiatry. 2018;17.

13. Davey CG, Pujol J, Harrison BJ. Mapping the self in the brain’s default mode network. NeuroImage. 2016;132.

14. Sui J, Gu X. Self as Object: Emerging Trends in Self Research. Trends in Neurosciences. 2017;40.

15. Dixon ML, Gross JJ. Dynamic network organization of the self: implications for affective experience. Current Opinion in Behavioral Sciences. 2021;39.

16. Halassa MM, Sherman SM. Thalamocortical Circuit Motifs: A General Framework. Neuron. 2019.

17. Bell PT, Shine JM. Subcortical contributions to large-scale network communication. Neuroscience and Biobehavioral Reviews. 2016;71.

18. Parnaudeau S, Bolkan SS, Kellendonk C. The Mediodorsal Thalamus: An Essential Partner of the Prefrontal Cortex for Cognition. Biological Psychiatry. 2018.

19. Pergola G, Danet L, Pitel AL, Carlesimo GA, Segobin S, Pariente J, et al. The Regulatory Role of the Human Mediodorsal Thalamus. Trends in Cognitive Sciences. 2018;22.

20. van der Werf YD, Scheltens P, Lindeboom J, Witter MP, Uylings HBM, Jolles J. Deficits of memory, executive functioning and attention following infarction in the thalamus; a study of 22 cases with localised lesions. Neuropsychologia. 2003;41.

21. Halassa MM, Kastner S. Thalamic functions in distributed cognitive control. Nature Neuroscience. 2017;20:1669–1679.

22. Greene DJ, Marek S, Gordon EM, Siegel JS, Gratton C, Laumann TO, et al. Integrative and Network-Specific Connectivity of the Basal Ganglia and Thalamus Defined in Individuals. Neuron. 2020;105.

23. Wolff M, Vann SD. The cognitive thalamus as a gateway to mental representations. Journal of Neuroscience. 2019;39.

24. Sheehan D v., Lecrubier Y, Sheehan KH, Janavs J, Weiller E, Keskiner A, et al. The validity of the Mini International Neuropsychiatric Interview (MINI) according to the SCID-P and its reliability. European Psychiatry. 1997;12.

25. Ehring T, Zetsche U, Weidacker K, Wahl K, Schönfeld S, Ehlers A. The Perseverative Thinking Questionnaire (PTQ): Validation of a content-independent measure of repetitive negative thinking. Journal of Behavior Therapy and Experimental Psychiatry. 2011;42.

26. Olszowy W, Aston J, Rua C, Williams GB. Accurate autocorrelation modeling substantially improves fMRI reliability. Nature Communications. 2019;10.

27. Friston KJ, Preller KH, Mathys C, Cagnan H, Heinzle J, Razi A, et al. Dynamic causal modelling revisited. NeuroImage. 2019;199.

28. Tak S, Noh J, Cheong C, Zeidman P, Razi A, Penny WD, et al. A validation of dynamic causal modelling for 7T fMRI. Journal of Neuroscience Methods. 2018;305.

29. Zeidman P, Jafarian A, Corbin N, Seghier ML, Razi A, Price CJ, et al. A guide to group effective connectivity analysis, part 1: First level analysis with DCM for fMRI. NeuroImage. 2019;200.

30. Zeidman P, Jafarian A, Seghier ML, Litvak V, Cagnan H, Price CJ, et al. A guide to group effective connectivity analysis, part 2: Second level analysis with PEB. NeuroImage. 2019;200.

31. Friston KJ, Litvak V, Oswal A, Razi A, Stephan KE, van Wijk BCM, et al. Bayesian model reduction and empirical Bayes for group (DCM) studies. NeuroImage. 2016;128.

32. Eckert U, Metzger CD, Buchmann JE, Kaufmann J, Osoba A, Li M, et al. Preferential networks of the mediodorsal nucleus and centromedian-parafascicular complex of the thalamus-A DTI tractography study. Human Brain Mapping. 2012;33.

33. van Kesteren MTR, Ruiter DJ, Fernández G, Henson RN. How schema and novelty augment memory formation. Trends in Neurosciences. 2012;35.

34. Denny BT, Kober H, Wager TD, Ochsner KN. A meta-analysis of functional neuroimaging studies of self- and other judgments reveals a spatial gradient for mentalizing in medial prefrontal cortex. Journal of Cognitive Neuroscience. 2012;24.

35. Dixon ML, de La Vega A, Mills C, Andrews-Hanna J, Spreng RN, Cole MW, et al. Heterogeneity within the frontoparietal control network and its relationship to the default and dorsal attention networks. Proceedings of the National Academy of Sciences of the United States of America. 2018;115.

36. Harrison BJ, Pujol J, Contreras-Rodríguez O, Soriano-Mas C, López-Solà M, Deus J, et al. Task-Induced deactivation from rest extends beyond the default mode brain network. PLoS ONE. 2011;6.

37. Alves PN, Foulon C, Karolis V, Bzdok D, Margulies DS, Volle E, et al. An improved neuroanatomical model of the default-mode network reconciles previous neuroimaging and neuropathological findings. Communications Biology. 2019. 2019. https://doi.org/10.1038/s42003-019-0611-3.

38. Schmitt LI, Wimmer RD, Nakajima M, Happ M, Mofakham S, Halassa MM. Thalamic amplification of cortical connectivity sustains attentional control. Nature. 2017. 2017. https://doi.org/10.1038/nature22073.

39. Saund J, Dautan D, Rostron C, Urcelay GP, Gerdjikov T v. Thalamic inputs to dorsomedial striatum are involved in inhibitory control: evidence from the five-choice serial reaction time task in rats. Psychopharmacology. 2017;234.

40. Rikhye R v., Wimmer RD, Halassa MM. Toward an Integrative Theory of Thalamic Function. Annual Review of Neuroscience. 2018;41.

41. Phillips JM, Fish LR, Kambi NA, Redinbaugh MJ, Mohanta S, Kecskemeti SR, et al. Topographic organization of connections between prefrontal cortex and mediodorsal thalamus: Evidence for a general principle of indirect thalamic pathways between directly connected cortical areas. NeuroImage. 2019;189.

42. Dehghani N, Wimmer RD. A computational perspective of the role of the thalamus in cognition. Neural Computation. 2019.

43. de Kloet SF, Bruinsma B, Terra H, Heistek TS, Passchier EMJ, van den Berg AR, et al. Bi-directional regulation of cognitive control by distinct prefrontal cortical output neurons to thalamus and striatum. Nature Communications. 2021;12.

44. Guo Z v., Inagaki HK, Daie K, Druckmann S, Gerfen CR, Svoboda K. Maintenance of persistent activity in a frontal thalamocortical loop. Nature. 2017;545.

45. Cash RFH, Weigand A, Zalesky A, Siddiqi SH, Downar J, Fitzgerald PB, et al. Using Brain Imaging to Improve Spatial Targeting of Transcranial Magnetic Stimulation for Depression. Biological Psychiatry. 2020.

