## Supplementary Information for "A thalamo-centric neural signature for restructuring negative self-beliefs"

### MRI Paradigm

Before scanning, participants were given detailed training on cognitive restructuring skills using Socratic questioning techniques, such as active rebuttal, reinterpretation, and perspective shifting [1]. After being shown a negative self-belief statement, participants were instructed to recall previous instances when the statement was not true or to imagine themselves as lawyers who must argue against the statement using factual information. To confirm participants' ability to carry out cognitive restructuring strategies, participants were asked to verbalize how they intended to mentally reframe example negative statements during scanning. Training was completed by a Masters-level clinical psychology trainee.

Prior to and following scanning, participants completed a 7-point self-rated questionnaire including the sixteen negative statements presented during the cognitive-restructuring task. Participants were instructed to indicate their level of agreement (1 = strongly disagree, 7 = strongly agree) with each statement. These statements were adapted from common negative self-beliefs described in CBT literature [1, 2], including statements such as: "I sometimes feel incompetent in the things I do" and "I think that I'm a failure". To verify the effectiveness of cognitive restructuring during the task, participant endorsement levels of negative self-beliefs pre- and post-scanning were compared using a paired sample t-test.

### Image Acquisition

Imaging was performed on a 7-Tesla (7T) research scanner (Siemens Healthcare, Erlangen, Germany) equipped with a 32-channel head-coil (Nova Medical Inc., Wilmington MA, USA). The functional sequence consisted of a multi-band (factor = 6) and grappa (2 times) accelerated GE-EPI sequence in the steady state (repetition time, 800ms; echo time, 22.2 ms; and pulse/flip angle, 45°) in a 20.8 cm field-of-view, with a 130 x 130-pixel matrix and a slice thickness of 1.6 mm (no gap; [3]. Eighty-four interleaved slices were acquired parallel to the anterior–posterior commissure line. A denoised T1-weighted high-resolution anatomical image (MP2RAGE; [3] was acquired for each participant to assist with functional time series co-registration (224 contiguous sagittal slices; repetition time = 5 seconds, echo time = 3.06 ms, flip angle=13°; in a 24 cm field of view, with a 256 x 256–pixel matrix and a slice thickness of 0.73 mm). To assist with head immobility, foam-padding inserts were placed either side of the participants' head. Cardiac and respiratory recordings were sampled at 50 Hz using a pulse-oximeter and respiratory belt. Information derived from these recordings were used for physiological noise correction (see below).

### Image Preprocessing and Physiological Noise Correction

Imaging data was pre-processed using Statistical Parametric Mapping (SPM) 12 (v7771, Welcome Trust Centre for Neuroimaging, London, UK) within a MATLAB 2019b environment (The

MathWorks Inc., Natick, MA). Motion artifacts were corrected by realigning each participant's time series to the mean image, and all images were resampled using 4th Degree B-Spline interpolation. Individualized motion regressors were created using the Motion Fingerprint approach to account for movement [4]. One participant was excluded due to a mean total scan-to-scan displacement over 1.6 mm (i.e., the size of one voxel). Each participant's anatomical images were co-registered to their respective mean functional image, segmented and normalized to the International Consortium of Brain Mapping template using the unified segmentation plus DARTEL approach. Smoothing was applied with a 3.2 $mm^3$ full-width-at-half-maximum (FWHM) gaussian kernel to preserve spatial specificity.

Physiological noise was modelled at the first level using the PhysIO Toolbox [5]. This toolbox applies noise correction to fMRI sequences using physiological recordings and has been found to improve blood-oxygen level dependent (BOLD) signal sensitivity and temporal rignal-to-noise ratio (tSNR) at 7T [6]. The Retrospective Image-based Correction function (RETROICOR; [7] was applied to model the periodic effects of heartbeat and breathing on BOLD signals, using acquired cardiac/respiratory phase information. The Respiratory Response Function (RRF; [8] convolved with respiration volume per time (RVT) was used to model low frequency signal fluctuations, which arose from changes in breathing depth and rate. Heart rate variability (HRV) was convolved with a predefined Cardiac Response Function (CRF; [9] to account for BOLD variances due to heartrate-dependent changes in blood oxygenation. Individualized DARTEL tissue maps segmented from each participant's respective anatomical scan were used to apply aCompCor, which models negative BOLD signals using principal components derived from white matter and CSF [10]. F-maps displaying group level gains following physiological noise modeling are available in Supplementary Information Fig. S1.

**Dynamic Causal Modelling (DCM)**

VOIs were calculated using the first eigenvariate of all voxels showing significant activation ($p < 0.05$, uncorrected) within a sphere with a radius of 4 mm around the subject specific maxima, with maxima required to be no more than 8 mm from the group maxima. To ensure that voxels included in MD VOIs did not capture adjacent regions and in accordance with recent research examining thalamic connectivity [11], time series extraction from the MD seed was run after applying an inclusive MD mask (excluding any non-MD voxels). The 1x1x1 $mm^3$ resolution atlas featured in the MNI-standardized Automated Anatomical Labelling 3 (AAL3) toolbox [12] was used to classify the location of the MD and only voxels falling within the medial magnocellular and lateral parvocellular nuclei of the MD were included in each VOI. Participant timeseries were pre-whitened in order to reduce serial correlations, high-pass filtered, and nuisance effects not covered by the 'effects of interest' F-contrast were regressed from the timeseries (i.e., 'adjusted' to the F-contrast).

**Table S1. Descriptive Statistics**

| | Mean (%) | SD | Minimum | Maximum |
|---|---|---|---|---|
| **Sex (%)** | 42.90% Female | | | |
| **Race/ethnicity (%)** | 54.8% Caucasian | | | |
| | 40.5% Asian | | | |
| | 4.8% Other | | | |
| **Age (years)** | 24.67 | 4.65 | 18 | 36 |
| **Years of education** | 16.56 | 2.64 | 13 | 23 |
| **PTQ Total** | 25.31 | 8.14 | 4 | 43 |

*n=42; PTQ: Perseverative Thinking Questionnaire.*

**Table S2. Cognitive Reframing Task General Linear Model (GLM) Results during Chal>Rep**

| Region | x | y | z | T | Ke |
|---|---|---|---|---|---|
| Supp_Motor_Area_L | -8 | 13 | 69 | 7.85 | 5224 |
| Caudate_L | -18 | 0 | 18 | 7.42 | 2924 |
| Cerebelum_Crus1_R | 42 | -59 | -34 | 6.67 | 2530 |
| Temporal_Pole_Sup_L | -51 | 14 | -21 | 6.18 | 4800 |
| Cerebelum_9_R | 2 | -50 | -43 | 5.92 | 313 |
| Cerebelum_8_R | 34 | -64 | -54 | 5.58 | 348 |
| Temporal_Pole_Sup_R | 51 | 11 | -22 | 5.51 | 205 |
| SN_pc_L | -6 | -26 | -16 | 5.4 | 61 |
| Frontal_Mid_2_L | -37 | 5 | 61 | 5.35 | 1817 |
| Olfactory_L | -5 | 22 | -11 | 5.17 | 860 |
| Caudate_R | 19 | -3 | 16 | 5.14 | 518 |
| Cerebelum_6_L | -29 | -48 | -34 | 5.13 | 170 |
| Vermis_6 | 5 | -59 | -24 | 4.7 | 639 |
| Caudate_R | 19 | -24 | 21 | 4.69 | 51 |
| Frontal_Mid_2_L | -29 | 51 | 6 | 4.67 | 278 |
| Cerebelum_8_L | -30 | -56 | -50 | 4.66 | 93 |
| Vermis_9 | 2 | -53 | -30 | 4.5 | 50 |
| OFCpost_L | -32 | 27 | -16 | 4.48 | 43 |
| Cingulate_Mid_L | -3 | -14 | 37 | 4.43 | 60 |
| Cerebelum_Crus1_L | -42 | -59 | -26 | 4.41 | 48 |
| Cerebelum_9_R | 5 | -35 | -53 | 4.23 | 48 |
| ACC_sup_R | 13 | 18 | 26 | 4.2 | 72 |
| Angular_L | -46 | -64 | 29 | 4.19 | 266 |
| ParaHippocampal_L | -21 | -22 | -27 | 4.17 | 37 |
| Cerebelum_8_R | 40 | -50 | -50 | 4.14 | 56 |
| Temporal_Mid_L | -54 | -56 | 21 | 4.1 | 38 |
| Temporal_Sup_R | 45 | -32 | -3 | 3.95 | 40 |
| Frontal_Med_Orb_R | 11 | 46 | -6 | 3.87 | 60 |

| | | | | | |
|---|---|---|---|---|---|
| Insula_R | 37 | 13 | 5 | 3.85 | 46 |
| Temporal_Pole_Mid_L | -34 | 21 | -34 | 3.84 | 51 |
| Vermis_4_5 | 2 | -64 | -10 | 3.61 | 32 |

*All results set at a $P_{FDR}$ < .05 threshold. Regions defined using the Automated Anatomical Labelling Atlas 3 (AAL3; local maxima labelling, 1 mm voxel edge; Rolls et al., 2020). Coordinates in MNI space.*

**Figure S1. Effects of PhysIO Toolbox noise correction on group-level effects**

**Cardiac Regressors**

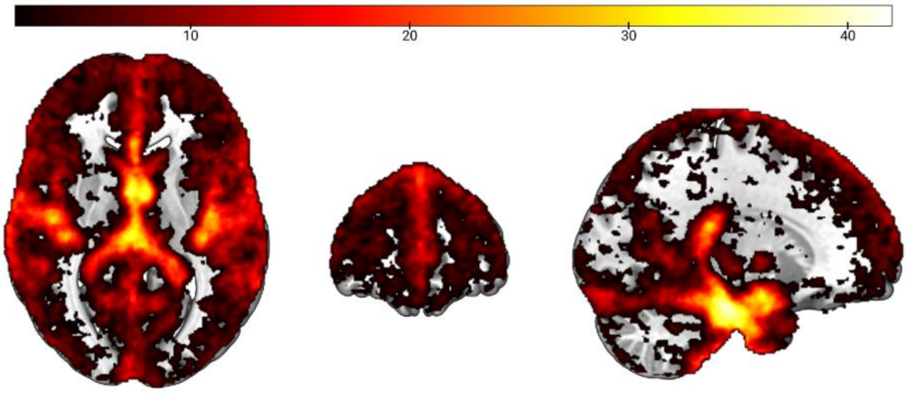


**Respiratory Regressors**

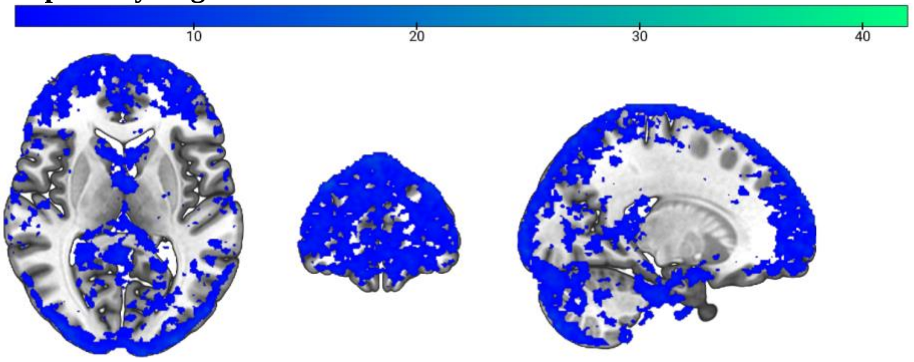


**Interaction Regressors (cardiac × respiration)**

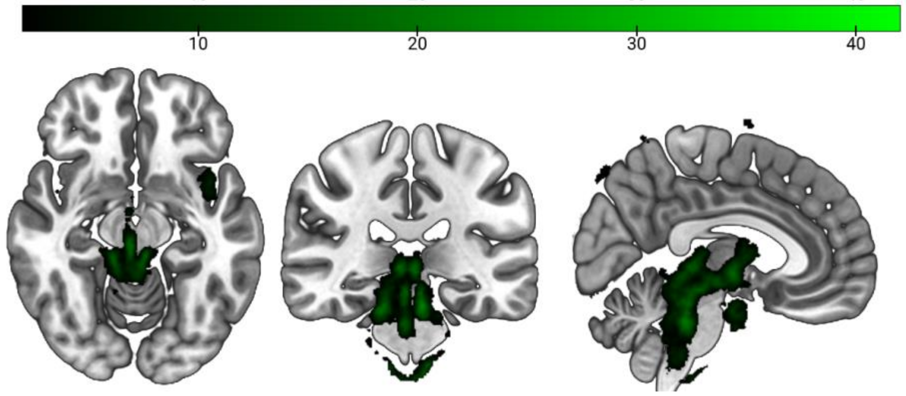


*Subject count of significant physiological noise correction using the PhysIO Toolbox (Kasper et al., 2017) for each voxel in individual 1st level analyses (peak level FWE-corrected p<0.05). The top row depicts the effects of cardiac regressors, the middle row depicts the effects of respiratory regressors, and the bottom row depicts the effects of interaction regressors (cardiac × respiration). The most prominent areas (brainstem, insula, thalamus) match physiological noise sites reported in the literature (Brooks et al., 2013, Hutton et al., 2011). Color bars represent voxel-wise subject-count.*

**Supplementary S2. Extrinsic between-region effective connectivity and its modulatory parameters during cognitive restructuring.**

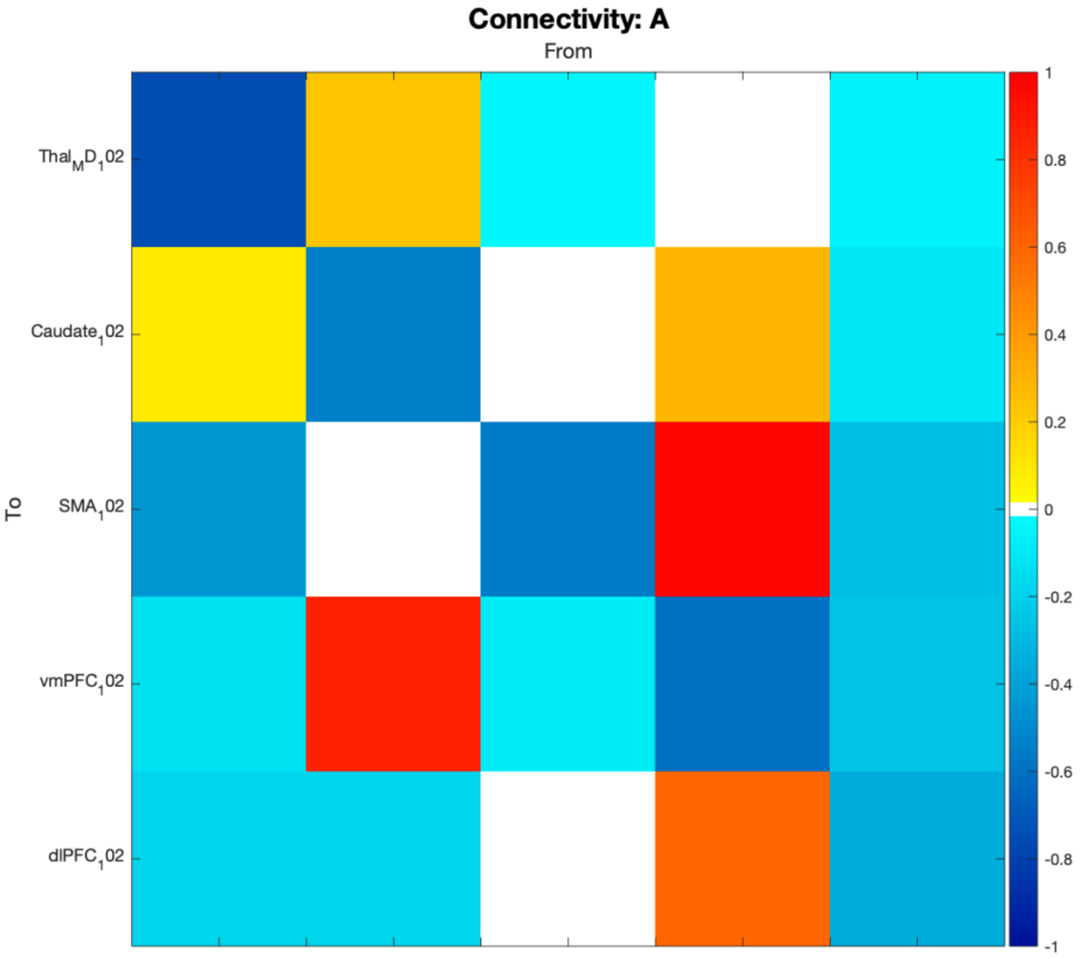

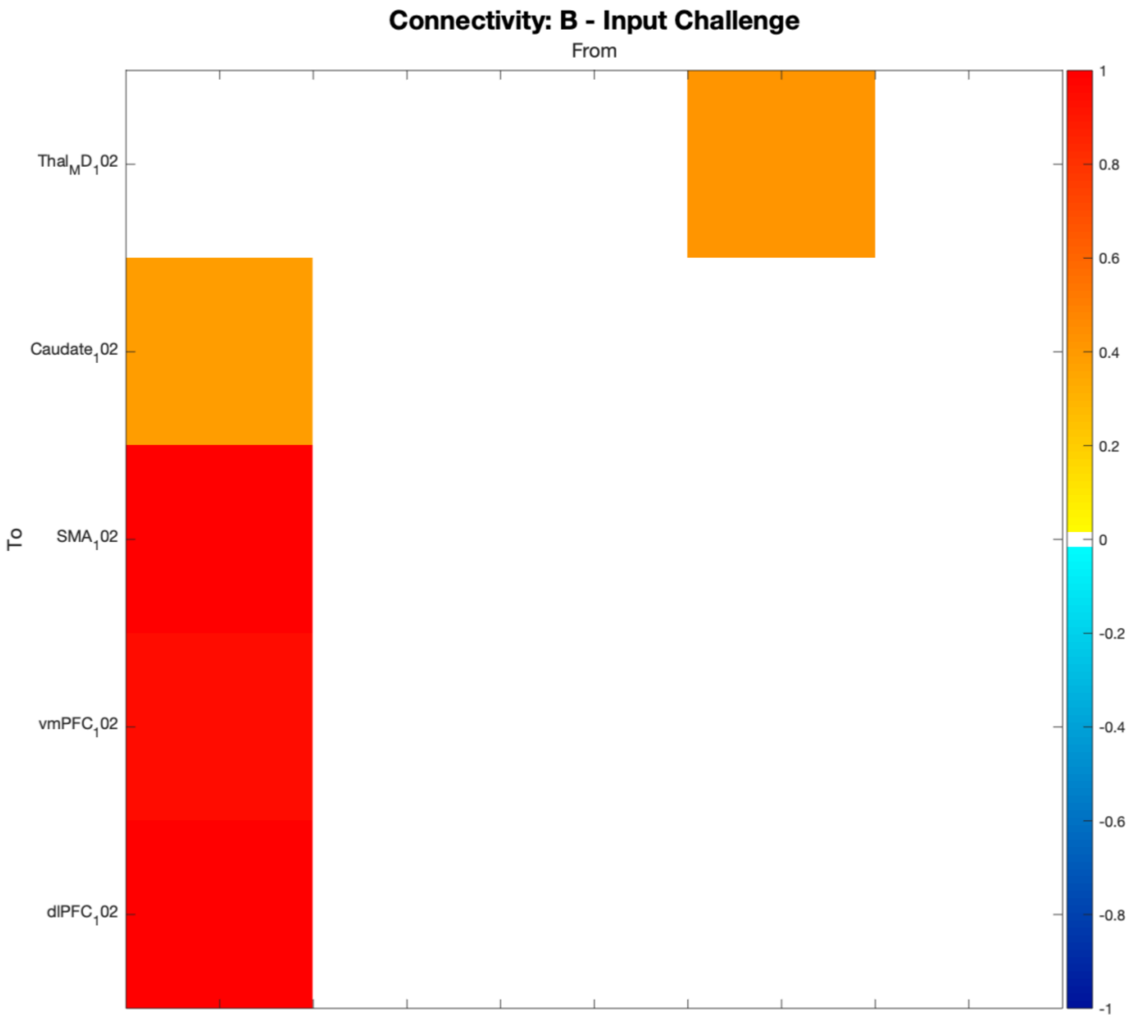


*Bayesian Model Averaging (BMA) of the PEB group-level effective connectivity models afforded the 'A' parameters representing baseline effective connectivity. BMA also yielded the modulatory 'B' parameters during cognitive structuring. The units of these effective connectivity and modulatory strengths are reported in Hertz. This figure only includes parameters related to connectivity with a posterior probability at or above 95 %. MD = mediodorsal thalamus; preSMA = presupplementary motor area; vmPFC = ventromedial prefrontal cortex; dlPFC = dorsolateral prefrontal cortex.*

**References**


1. Beck AT, Dozois DJA. Cognitive therapy: Current status and future directions. Annual Review of Medicine. 2011;62.
2. Osmo F, Duran V, Wenzel A, de Oliveira IR, Nepomuceno S, Madeira M, et al. The negative core beliefs inventory: Development and psychometric properties. Journal of Cognitive Psychotherapy. 2018;32.
3. Setsompop K, Gagoski BA, Polimeni JR, Witzel T, Wedeen VJ, Wald LL. Blipped-controlled aliasing in parallel imaging for simultaneous multislice echo planar imaging with reduced g-factor penalty. Magnetic Resonance in Medicine. 2012;67.
4. Wilke M. An alternative approach towards assessing and accounting for individual motion in fMRI timeseries. NeuroImage. 2012;59.
5. Kasper L, Bollmann S, Diaconescu AO, Hutton C, Heinzle J, Iglesias S, et al. The PhysIO Toolbox for Modeling Physiological Noise in fMRI Data. Journal of Neuroscience Methods. 2017;276.
6. Reynaud O, Jorge J, Gruetter R, Marques JP, van der Zwaag W. Influence of physiological noise on accelerated 2D and 3D resting state functional MRI data at 7 T. Magnetic Resonance in Medicine. 2017;78.
7. Glover GH, Li TQ, Ress D. Image-based method for retrospective correction of physiological motion effects in fMRI: RETROICOR. Magnetic Resonance in Medicine. 2000;44.
8. Birn RM, Smith MA, Jones TB, Bandettini PA. The respiration response function: The temporal dynamics of fMRI signal fluctuations related to changes in respiration. NeuroImage. 2008;40.
9. Chang C, Cunningham JP, Glover GH. Influence of heart rate on the BOLD signal: The cardiac response function. NeuroImage. 2009;44.
10. Behzadi Y, Restom K, Liau J, Liu TT. A component based noise correction method (CompCor) for BOLD and perfusion based fMRI. NeuroImage. 2007;37.
11. Kafkas A, Mayes AR, Montaldi D. Thalamic-Medial Temporal Lobe Connectivity Underpins Familiarity Memory. Cerebral Cortex. 2020. 2020. https://doi.org/10.1093/cercor/bhz345.
12. Rolls ET, Huang CC, Lin CP, Feng J, Joliot M. Automated anatomical labelling atlas 3. NeuroImage. 2020;206.
